# Development of the visual white matter pathways mediates development of electrophysiological responses in visual cortex

**DOI:** 10.1101/2021.05.26.445879

**Authors:** Sendy Caffarra, Sung Jun Joo, David Bloom, John Kruper, Ariel Rokem, Jason D. Yeatman

## Abstract

The latency of neural responses in the visual cortex changes systematically across the lifespan. Here we test the hypothesis that development of visual white matter pathways mediates maturational changes in the latency of visual signals. Thirty-eight children participated in a cross-sectional study including diffusion MRI and MEG sessions. During the MEG acquisition, participants performed a lexical decision and a fixation task on words presented at varying levels of contrast and noise. For all stimuli and tasks, early evoked fields were observed around 100 ms after stimulus onset (M100), with slower and lower amplitude responses for low as compared to high contrast stimuli. The optic radiations and optic tracts were identified in each individual’s brain based on diffusion MRI tractography. The diffusion properties of the optic radiations predicted M100 responses, especially for high contrast stimuli. Higher optic radiation fractional anisotropy (FA) values were associated with faster and larger M100 responses. Over this developmental window, the M100 responses to high contrast stimuli became faster with age and the optic radiation FA mediated this effect. These findings suggest that the maturation of the optic radiations over childhood accounts for individual variations observed in the developmental trajectory of visual cortex responses.

## Introduction

Electrophysiological responses in visual cortex are subject to a high degree of variability. It is known that these responses reliably differ among individuals and change over development (Allison et al. 1984; Onofrj et al. 2001). Being able to account for these variations will improve our understanding of the brain circuits and their developmental trajectories in health and disease. One possible source of variability in electrophysiology lies in structural properties of the white matter tracts (Kanai and Rees 2011; Wandell 2016), which carry signals to the cortex. In the last two decades, novel neuroscientific tools (including tractography) have opened the possibility to address this question and explore the relation between functional and structural properties of the brain (Jeurissen et al. 2019; Wandell 2016). In this study we combine diffusion magnetic resonance imaging (dMRI) and tractography with magnetoencephalography (MEG), to examine the variability of visual responses during childhood. We asked whether developmental differences in visual response properties are the result of the maturation of the white matter pathways carrying visual signals. Specifically, we test the hypothesis that developmental variations of visual pathways mediate age effects on electrophysiological responses.

In neurologically healthy individuals, any visual input elicits electrophysiological responses roughly 100 ms after stimulus presentation, though the precise timing varies by as much as 50 ms among individuals (Allison et al. 1984; Kolb, Fernandez, and Nelson 2005; Odom et al. 2004; Spear 1993; Vialatte et al. 2010). These visually evoked responses can be recorded over the occipital part of the scalp and have their neural source in the early visual cortex. There is consistent evidence that the latency of visual responses change substantially across the lifespan, with a speed increase of 10 ms per decade within the first 20 years of age and a symmetrical slow down after 60 years of age (Armstrong, Slaven, and Harding 1991; Allison et al. 1984; Onofrj et al. 2001; Spear 1993). The nature of these latency changes is still poorly understood, but clinical research suggests that the structural properties of visual white matter pathways can play a crucial role in determining the latency of visual signals. Studies on patients with demyelinating lesions of the visual tracts have shown that white matter diffusion properties (such as fractional anisotropy, FA, or mean diffusivity, MD) predict delays in the electrophysiological responses of the visual cortex (Alshowaeir et al. 2014; Berman et al. 2020; Kolbe et al. 2012; Lobsien et al. 2014; Naismith et al. 2010; M. Y. Takemura et al. 2017). For instance, patients with lower FA or higher MD of the optic radiations (the tract connecting the lateral geniculate nucleus to the primary visual cortex; Figure 1) showed slower visually evoked responses. These results suggest that the demyelination of visual pathways (reflected by altered diffusion properties) accounts for conduction delays of visual signals that are carried from the eyes to the visual cortex, and ultimately explains the latency variability of evoked responses recorded on the scalp in patient populations.

**Figure 1.**
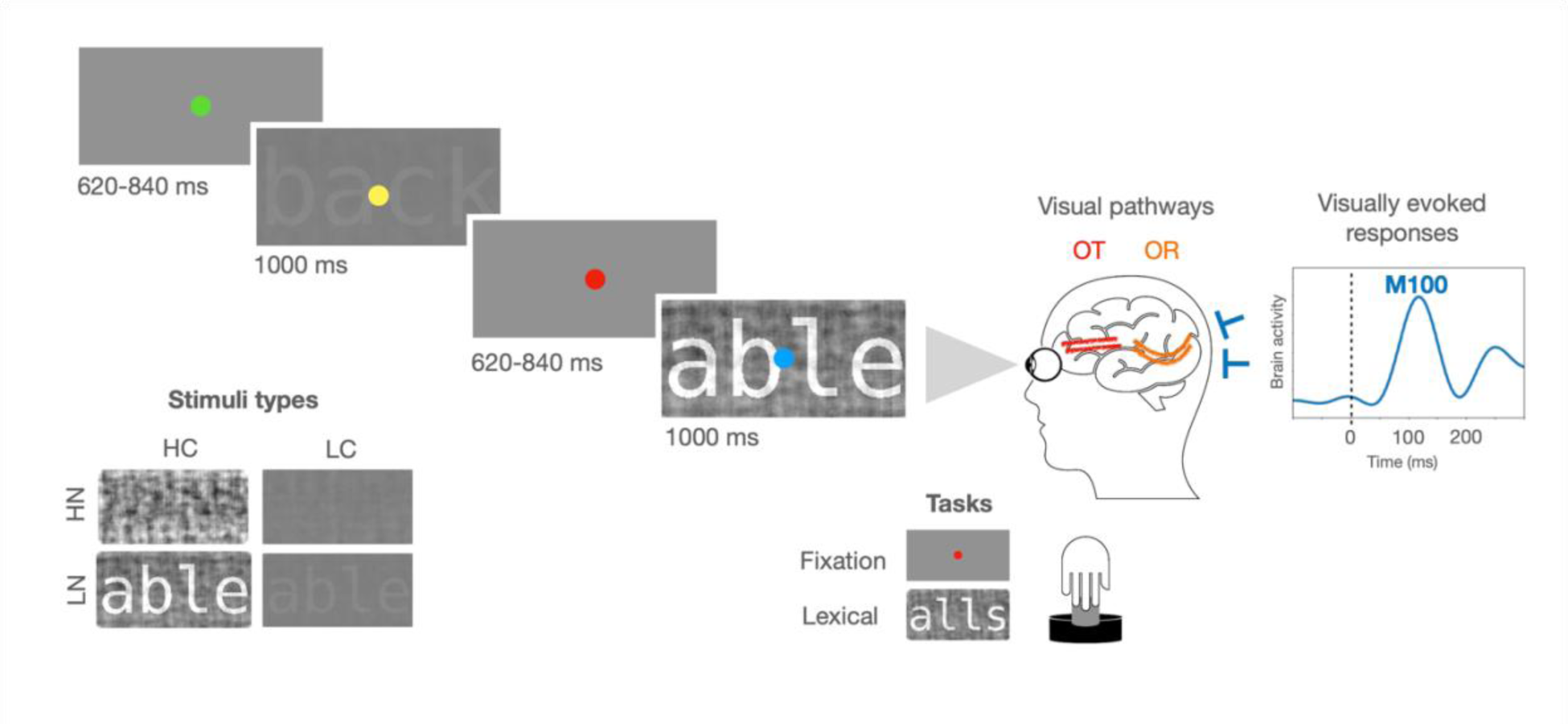
Schematic representation of an MEG experimental trial, the experimental conditions, and the visual pathways. On each trial, the stimulus was shown at a high or low level of contrast (HC or LC), and at a high or low degree of noise (HN or LN). The task required a button press whenever the fixation dot turned red (fixation task) or whenever a pseudoword was presented (lexical decision task). Visual information is received by the eyes and carried through the visual pathways (including the optic nerve, the optic tract (OT) and the optic radiation (OR)) to the visual cortex, where visually evoked responses can be recorded through MEG sensors. A magnetic evoked response peaking around 100 ms (M100 response, dotted line) from a participant in the present study (S172: male, 9 years old) is displayed here as a representative example of visually evoked response.

The relation between visual white matter properties and electrophysiology has recently been reported in one study of healthy adults (H. Takemura, Yuasa, and Amano 2020). This suggests that there may be sufficient variability in the organization of the optic radiations among typical adults that the structural differences affect signaling properties in the visual system. Moreover, three additional studies have reported a similar structural-functional relationship in the infant (1-4 months, Dubois et al. 2008; 1-6 months, Adibpour, Dubois, and Dehaene-Lambertz 2018) and in the ageing brain (18-88 years, (Price et al. 2017). Despite these new findings, the amount of evidence showing links between electrophysiological responses and white matter properties is still scarce, and mainly related to clinical populations. In addition, the age range from 1 to 18 years old remains fully unexplored.

Childhood represents a crucial developmental phase where many visual circuits reach maturity (Allison et al. 1984; Gilmore, Knickmeyer, and Gao 2018; Onofrj et al. 2001; Siu and Murphy 2018). Moreover, this is a developmental window where white matter pathways are still rapidly changing and approaching maturity (Lebel et al. 2008, 2012; Yeatman, Wandell, and Mezer 2014). However, the potential relationship between these structural and functional maturational variations is yet to be described. The present study links the structural and functional properties of the visual system in childhood by combining dMRI with MEG. We show that the maturational differences of the optic radiations observed during childhood mediate age effects on visually evoked responses.

## Materials and Methods

### Participants

Forty-six children participated in a cross-sectional study including a dMRI and an MEG session. All the data were manually inspected for quality of the MEG recordings and artifacts in the dMRI data, and 8 participants were excluded leaving a final sample size of thirty-eight (17 females, mean age: 9.5 y, SD: 1.6, age range: 7-12 y, between-sessions time gap: 0-35 dd; only two participants had a time gap different from zero and this time difference did not affect the main results reported here, see Supplementary Materials S3). All participants had normal or corrected-to-normal vision, and no history of neurological, psychiatric, or sensory disorder. All parents (or legal guardians) signed a written informed consent, and all participants gave their verbal assent. The study was conducted in accordance with the guidelines of the University of Washington Subjects Division and was approved by the Institutional Review Board. Participant recruitment met Human Brain Mapping expectation of inclusivity other than as required scientifically.

Based on the average of the correlations reported in a previous study of healthy adults using a similar methodology (*r*avg=0.48; H. Takemura, Yuasa, and Amano 2020), the present sample size ensures a statistical power of at least 0.87 (Hulley et al. 2001).

### MEG materials and experimental design

The MEG data analyzed here came from a previous study on automaticity in the brain’s reading circuitry (Joo et al. 2021). In the present study we focused on **early** MEG responses evoked by the visual stimuli. The M100 response was the focus of our analysis given its large amplitude and its high signal-to-noise ratio (Allison et al. 1984; Onofrj et al. 2001), which makes it a prominent early visual response that can be reliably measured in children. Visual words were presented at two contrast levels (high and low: HC and LC) and two noise levels (high and low: HN and LN). Two hundred and forty images of four-letter English words were rendered in Courier font. Visual stimuli had a Weber contrast of 7.8% or 100%. In addition, each visual stimulus was mixed with a different percentage of noise (20% or 80%), corresponding to the phase-scrambled version of the original image. This procedure led to the creation of readable (20% of noise, LN) and unreadable experimental stimuli (80% of noise, HN).

The images were presented in four separate runs and repeated twice (eight runs in total; 60 experimental stimuli per run). Two different tasks were carried out in alternating runs on the identical set of stimuli: a fixation task and a lexical decision task. In order to make these tasks possible, a colored fixation dot was added at the center of the screen and a small set of pseudowords (n=11) was presented for each run together with the rest of the experimental stimuli. In the fixation task, participants had to press a response button when a fixation dot turned red. In the lexical decision task, participants had to press a response button when a pseudoword was presented.

During each MEG experimental trial, a visual stimulus appeared on the screen for 1 sec and was followed by a blank with a random duration between 620 and 840 ms (see Figure 1). A colored fixation dot was always present in the center of the screen and changed its color every 500 ms (among the following options: green, blue, yellow, cyan or red). The stimuli were presented on a gray background (50 cd/m^2^) and subtended 2.7 degrees at a viewing distance of 1.25 m.

### MEG acquisition and pre-processing

MEG data were recorded in a magnetically shielded room (Maxshieldł, Elekta Oy, Helsinki, Finland) using an Elekta-Neuromag MEG device (including 102 sensors with two planar gradiometers and one magnetometer each). MEG recordings were acquired continuously with children in sitting position, with a bandpass filter at 0.01−600 Hz and a sampling rate of 1.2 KHz. Head position inside the helmet was continuously monitored using head position indicator coils. The location of each coil relative to the anatomical fiducials (nasion, and left and right preauricular points) was defined with a 3D digitizer (Polhemus Fastrak, Colchester, VT, USA). About 100 head surface points were digitized.

MEG data were analyzed using MNE-Python (Gramfort et al. 2013). The signal was subjected to noise reduction using the Maxwell filter function and data were individually corrected for head movements using the average of participants’ initial head positions as a reference. The temporally extended signal space separation method was applied with a correlation limit of 0.98 and a segment length of 10 sec (Taulu and Hari 2009; Taulu and Kajola 2005). Bad channels were substituted with interpolated values. An Independent Component Analysis (ICA) was applied to the down-sampled and filtered MEG continuous signal (each 5th data point was selected, Gramfort et al. 2013). Downsampling was only used for the ICA analysis and it was not applied in the subsequent preprocessing steps. Independent components corresponding to the heartbeat and ocular artifacts were automatically identified and removed from the filtered MEG signal based on cross-trial phase statistics with the ECG and EOG channels (Dammers et al. 2008). The average number of rejected components was 3 (SD: 1.7). MEG epochs were obtained, including 0.6 sec before and 1.6 second after the visual presentation onset. Residual artifacts exceeding a peak-to-peak amplitude of 1000e-12 fT/cm for gradiometers and of 4000e-14 fT for magnetometers were automatically rejected. On average, 6% (SD: 8.4) of trials were rejected, with no significant difference across conditions (*F*(7,296)=0.09, *p*=0.99).

To obtain evoked related fields (ERFs), artifact-free trials were low-pass filtered (firwin method was used with upper passband edge of 40 Hz, filter length: 331 ms), averaged and baseline corrected (-0.6-0 s). ERFs were calculated for each condition and each participant and they were quantified by computing the root mean square of the two gradiometers in each pair.

For each main effect (contrast, noise, and task; 240 trials per condition), ERFs were statistically compared using a nonparametric cluster-based permutation test (Maris and Oostenveld 2007). Specifically, *t*-statistics were computed for each sensor and time point during the 0 – 800 ms time window, and a clustering algorithm formed groups of channels over time points based on these tests. In order for a data point to become part of a cluster, a threshold of p = 0.05 was used (based on a two-tailed t-test, only vertices with data values more extreme than t>8 were included in the cluster). The sum of the t-statistics in a sensor group was then used as a cluster-level statistic, which was then tested with a randomization test using 1000 runs.

The M100 peak was identified for each participant as the highest amplitude fluctuation within the spatio-temporal cluster previously identified by the cluster-based permutation (8 occipital sensors, 80-150 ms; see Figure 2). The M100 latency was estimated as the time point at which the M100 amplitude reached 50% of the peak value (this relative criterion represents one of the most accurate methods to estimate the latency of evoked responses and it ensures higher statistical power compared to absolute criteria, Kiesel et al. 2008). The M100 amplitude at the half peak latency was also extracted for each participant.

**Figure 2.**
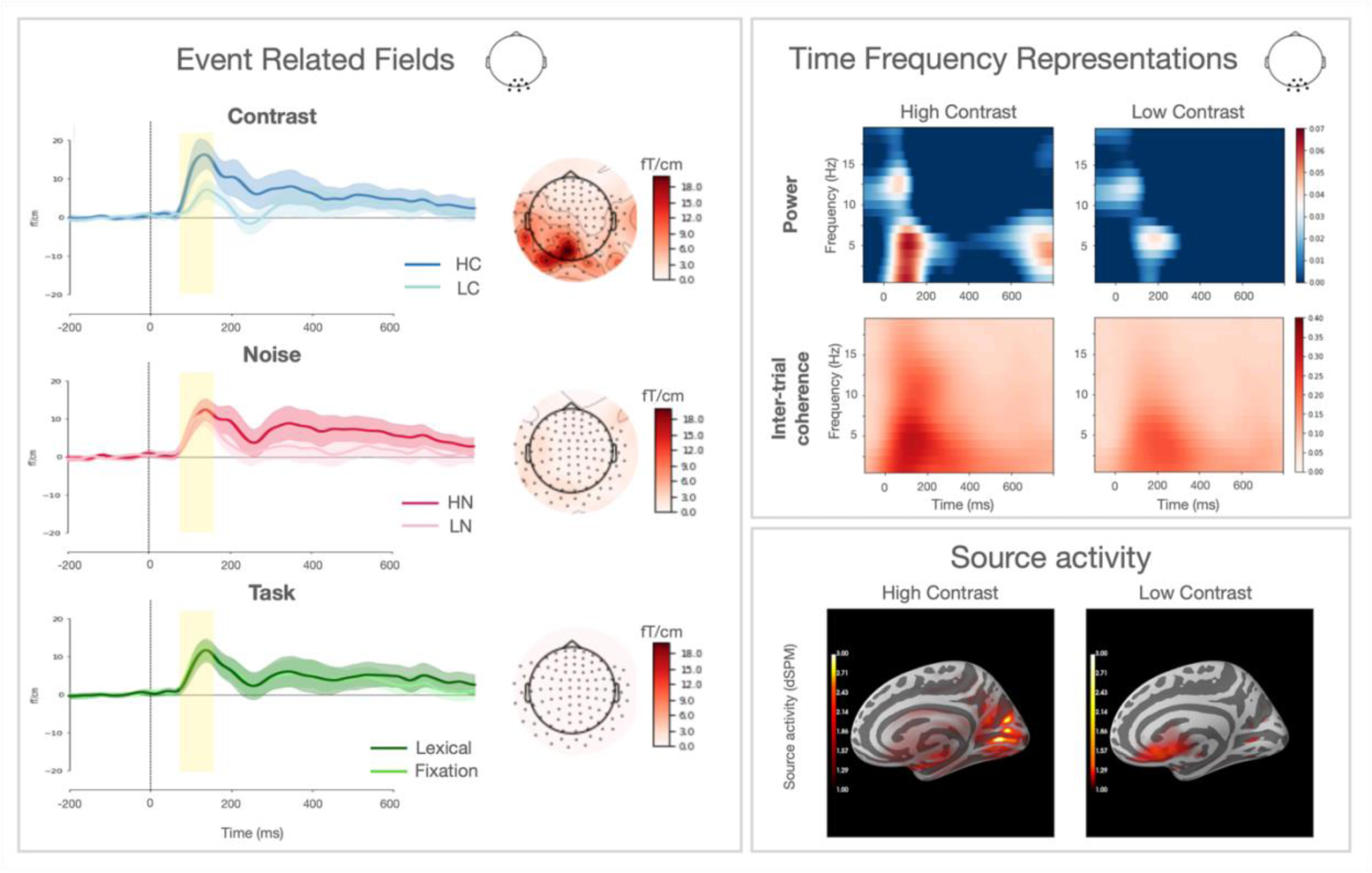
**ERFs panel:** ERFs responses to different levels of contrast, noise, and task over an occipital cluster of sensors (displayed beside the title panel). The yellow box represents the time window where the main effect of contrast reached its maximum (80-150 ms). Topographic distribution of each effect at 100 ms (calculated as the difference within each condition pair) is displayed on the right side. **TFRs panel:** TFRs of power and inter-trial coherence for occipital sensors. Only HC and LC conditions are represented here. Power values are expressed as the relative change (logratio) from a baseline interval between -0.4 and -0.2 ms. **Source Activity panel:** the neural source of the M100 response is represented for the HC and LC conditions.

Any significant ERFs effect was reconstructed in the source space. A cortical MEG source space was constructed using dipoles with 3 mm spacing. These were constrained to be normal to the cortical surface using Freesurfer (Dale, Fischl, and Sereno 1999). A forward solution was calculated to map dipole currents in the source space to the sensor space (Mosher et al. 1999). Dipole currents in this whole-brain source space were estimated from the evoked MEG response. A minimum-norm linear estimation (MNE) approach was employed (Dale, Fischl, and Sereno 1999; Hämäläinen and Ilmoniemi 1994; Hämäläinen and Sarvas 1989) with sensor noise covariance estimated from 100 ms epochs prior to each trial onset. Source localization data were then mapped to an average brain (freesurfer averaged brain) using a non-linear spherical morphing procedure (20 smoothing steps) that optimally aligns individual sulcal–gyral patterns (Fischl et al. 1999).

To more completely describe the electrophysiological properties of visually evoked responses, we characterized the M100 response within the time-frequency domain. This method explores single MEG trials within a narrow frequency band. By doing this, the amplitude of the noise is considerably reduced as compared to single-trial evoked responses (Herrmann et al. 2014). Also, the interpretation of time-frequency properties can be better related to the degree of synchronization of neural oscillatory activity (Z. Zhang 2019).

Time-frequency representations (TFRs) of power and inter-trial coherence values were calculated using Morlet wavelets on the unfiltered data. Single-trial time-frequency values were decomposed between 1 and 20 Hz (in steps of 1Hz), with a width of the wavelets equal to the half of the frequency under interest. The resulting values were averaged across trials for each condition and each participant. Our visually evoked responses were evident also in the time-frequency domain as the most prominent power and inter-trial coherence modulation over occipital sensors around 100 ms after stimulus presentation (see Figure 2). Maximum values of M100 power and inter-trial coherence were extracted for each subject using a similar spatio-temporal cluster employed for the ERFs values extraction (8 occipital sensors, 0-200 ms, 1-8 Hz; this was the result of a cluster-based permutation analogous to the one performed for the ERFs).

### MRI acquisition and pre-processing

Imaging data were recorded using an 8-channel phased-array SENSE head coil on a 3 T Phillips Achieva scanner (Philips, Eindhoven, Netherlands) at the University of Washington Diagnostic Imaging Sciences Center with a 32-channel head coil. A whole-brain anatomical volume at 0.8 x 0.8 x 0.8 mm resolution was acquired using a T1- weighted MPRAGE sequence (TR 15.22 s, TE 3 ms, matrix size 320 x 320, field of view 240 x 256 x 169.6, 212 slices). Head motion was minimized by an inflated cap, and participants were monitored through a closed-circuit camera system. Diffusion-weighted magnetic resonance imaging (dMRI) data were acquired with a spatial resolution of 2.0 mm^3^ and full brain coverage (phase encoding direction: anterior-posterior). A diffusion-weighted imaging (DWI) scan was acquired with 64 non-collinear directions (b-value = 2000 s/mm^2^) with a TR of 7700 ms and a TE of 85 ms. The DWI scan also contained 4 volumes without diffusion weighting (b-value = 0). In addition, one scan with six non-diffusion-weighted volumes with a reversed phase encoding direction (posterior-anterior) was collected to correct for echo-planar imaging distortions related to inhomogeneities in the magnetic field.

The T1-weighted (T1w) images were corrected for intensity non-uniformity (INU) using N4BiasFieldCorrection (Tustison et al. 2010, ANTs 2.3.1), and used as T1w- reference throughout the workflow. The T1w-reference was then skull-stripped using antsBrainExtraction.sh (ANTs 2.3.1), using OASIS as target template. Spatial normalization to the ICBM 152 Nonlinear Asymmetrical template version 2009c (Fonov et al. 2009) was performed through nonlinear registration with antsRegistration (ANTs 2.3.1, (Avants et al. 2008), using brain-extracted versions of both T1w volume and template. Brain tissue segmentation of cerebrospinal fluid (CSF), white-matter (WM) and gray-matter (GM) was performed on the brain-extracted T1w using FAST (FSL 6.0.3, Y. Zhang, Brady, and Smith 2001).

Preprocessing and reconstruction were carried out using QSIprep 0.11.5 (https://qsiprep.readthedocs.io/, based on Nipype 1.5.0; Cieslak et al. 2020; K.

Gorgolewski et al. 2011; K. J. Gorgolewski et al. 2018), which included topup distortion, motion and eddy current correction (Andersson and Sotiropoulos 2016; Andersson, Skare, and Ashburner 2003; Smith et al. 2004). Multi-tissue fiber response functions were estimated using the dhollander algorithm as implemented in MRtrix3 (Tournier et al. 2019). Fiber orientation distributions (FODs) in each voxel were estimated via constrained spherical deconvolution (CSD, Tournier et al. 2004, 2008) using an unsupervised multi-tissue method (T. Dhollander et al. 2019; Thijs Dhollander, Raffelt, and Connelly 2016). FODs were intensity-normalized using mtnormalize (Raffelt et al. 2017). Probabilistic tractography was carried out using the default parameters implemented in QSIprep (10M streamlines, minimum length: 30 mm, maximum length: 250 mm). The left and the right optic radiations were identified using vistasoft (https://github.com/vistalab/vistasoft) based on two endpoint regions of interest (ROIs) corresponding to the primary visual cortex and the central part of the thalamus including the lateral geniculate nucleus (defined based on the AICHA atlas, Joliot et al. 2015; minimum distance 3 mm). To further clean the tract from crossing fibers (Fan et al. 2016; Sherbondy et al. 2008), three exclusion ROIs were also used (temporal pole, and occipital pole from the AICHA atlas, and the posterior portion of the thalamus based on the brainnetome atlas; minimum distance 3 mm, Fan et al. 2016; Sherbondy et al. 2008). All ROIs were defined in a MNI template and transformed to each participant’s native space using QSIprep. A final cleaning step was carried out to remove outlier fibers based on streamline average length and mean Gaussian distance from the bundle core (threshold of 3 SD, streamlines were resampled to 4 nodes during the outlier cleaning phase; (Yeatman et al. 2012). The optic radiations were clipped at the endpoint ROIs (minimum distance: 3 mm) to avoid potential partial volume effects at the white matter/gray matter border (see Supplementary Materials S1 for a schematic representation of the bundle segmentation pipeline). The diffusion data was then fitted with the tensor model using a standard least-squares algorithm. Diffusion metrics were projected onto the previously identified optic radiations and fractional anisotropy (FA) was mapped onto each tract. FA values along the tract were weighted based on each streamline’s distance from the core of the tract (Yeatman et al. 2012). For each participant, the FA values of the left and right optic radiations were averaged.

A similar approach was taken to identify the optic tract. A first endpoint ROI corresponded to the central part of the thalamus used for the optic radiations (minimum distance 3 mm). A second waypoint ROI (minimum distance 3 mm) was manually defined at the center of the optic chiasm of each participant (5 mm sphere; mean coordinates of the center: 0, +3, -16). The tract was further cleaned by removing outlier fibers based on streamline average length and mean distance from the bundle core (threshold of 2 and 3 SD, respectively). The average FA values of the left and right optic tracts were calculated for each participant.

Two additional white matter tracts that were not part of the visual system (left and right uncinate and corticospinal tract) were segmented and average FA values were calculated using pyAFQ (https://yeatmanlab.github.io/pyAFQ; Yeatman et al. 2012; Kruper et al. 2021). These tracts were used as control pathways as they do not carry visual input from retina to early visual cortex.

### Statistical analysis

Statistical analyses were carried out with R 4.0.0 (https://www.R-project.org/). To test the relationship between structural properties of the optic radiations and latency of visual responses, a Pearson correlation between mean FA values and M100 latency was calculated. A biweight midcorrelation was also added as a median-based measure of similarity that is less sensitive to outliers and can be used as a robust alternative to mean-based similarity estimates, such as Pearson correlation (Langfelder and Horvath 2012). To make sure that the FA effect on electrophysiology held after accounting for developmental differences, M100 latency was also analyzed using a linear regression model including FA and Age as predictors. The effect of age on electrophysiological responses was further examined using a causal mediation analysis (as implemented in (Tingley et al. 2014). Two linear regression models were initially specified: a mediator model estimating the effect of age on FA, and an outcome model estimating the effect of FA and age on electrophysiological responses. The fitted objects of these two models were the inputs to the mediate function of the mediation R package (Tingley et al. 2014), which computed the average causal mediation effect (indirect effect of age on electrophysiology that is related to the FA mediator) and the average direct effect (effect of age on electrophysiology after partialling out the effect of the FA mediator). The sum of these two effects resulted in the total effect of age on electrophysiology. A bootstrap using 1000 simulations was used to calculate the uncertainty estimates of these mediation results (Efron and Tibshirani 1994).

Secondary analyses concerned the other properties of the M100 response. Similar FA-M100 correlations, regressions and mediation analyses were carried out with additional electrophysiological properties of the M100 response (M100 peak amplitude, power, and inter-trial coherence) to further understand the relation between structural and functional properties of the visual brain network.

## Results

### Behavioral results

Participants paid attention to the visual stimuli, as shown by the intermediate-to-high levels of accuracy in the fixation task (median accuracy: 81% correct, IQR: 35; median RT: 547 ms, IQR: 124). The lexical task showed lower accuracy and slower responses as a result of children’s variable reading skills (median: 17%, IQR: 15; median RT: 880 ms, IQR: 391). No significant correlation was observed between these behavioral performances and the structural or functional brain measures described below (all *p*s>.05).

### MEG results

The M100 responses showed substantial modulations based on the contrast of the image (see Figure 2) and did not change based on image noise or cognitive task. In accordance with previous electrophysiological studies (Abdullah et al. 2012; Gebodh, Vanegas, and Kelly 2017; Maddess, James, and Bowman 2005), high contrast stimuli elicited larger and faster M100 responses as compared to low contrast stimuli (*p*<.001). The response was 8 ms faster and 57% larger for high contrast compared to low contrast stimuli. The M100 contrast effect was centro-posteriorly distributed (Figure 2, left panel) and its source was localized bilaterally in the early visual cortex (Figure 2, right panel). A precise hemispheric characterization of the M100 responses was not possible due to spatial leakage issues in MEG analysis (Hauk, Stenroos, and Treder 2019). For this reason, the M100 individual responses analyzed here are the result of the average between left and right sensors. No other main effects (noise or task) or interactions could be observed in the M100 response (*ps*>.05).

### Linking MEG responses to visual white matter pathways

High contrast stimuli elicited sharper and more reliable evoked responses in the visual cortex (reliability measures calculated by correlating individual MEG measures of two halves of randomized trials: high contrast stimuli r=0.70; low contrast stimuli r=0.56). For this reason, we mainly focus on the high contrast condition and results from the low contrast stimuli are reported in Supplementary Materials S2. Individual differences could be observed in M100 responses, with some participants reaching the maximum response amplitude at shorter latencies than others (Figure 3, left panel). Similarly, individual variability could be also appreciated in the optic radiations FA values (Figure 3, middle panel). Despite these FA variations, participants consistently showed higher FA values in the left optic radiation. This hemispheric difference is in line with previous literature reports (e.g., diffusivity 7% higher in the right hemisphere; (Dayan et al. 2015; Levin et al. 2010; Sherbondy et al. 2008; Xie et al. 2007) and may be related to the different patterns of crossing fibers in the two hemispheres.

**Figure 3.**
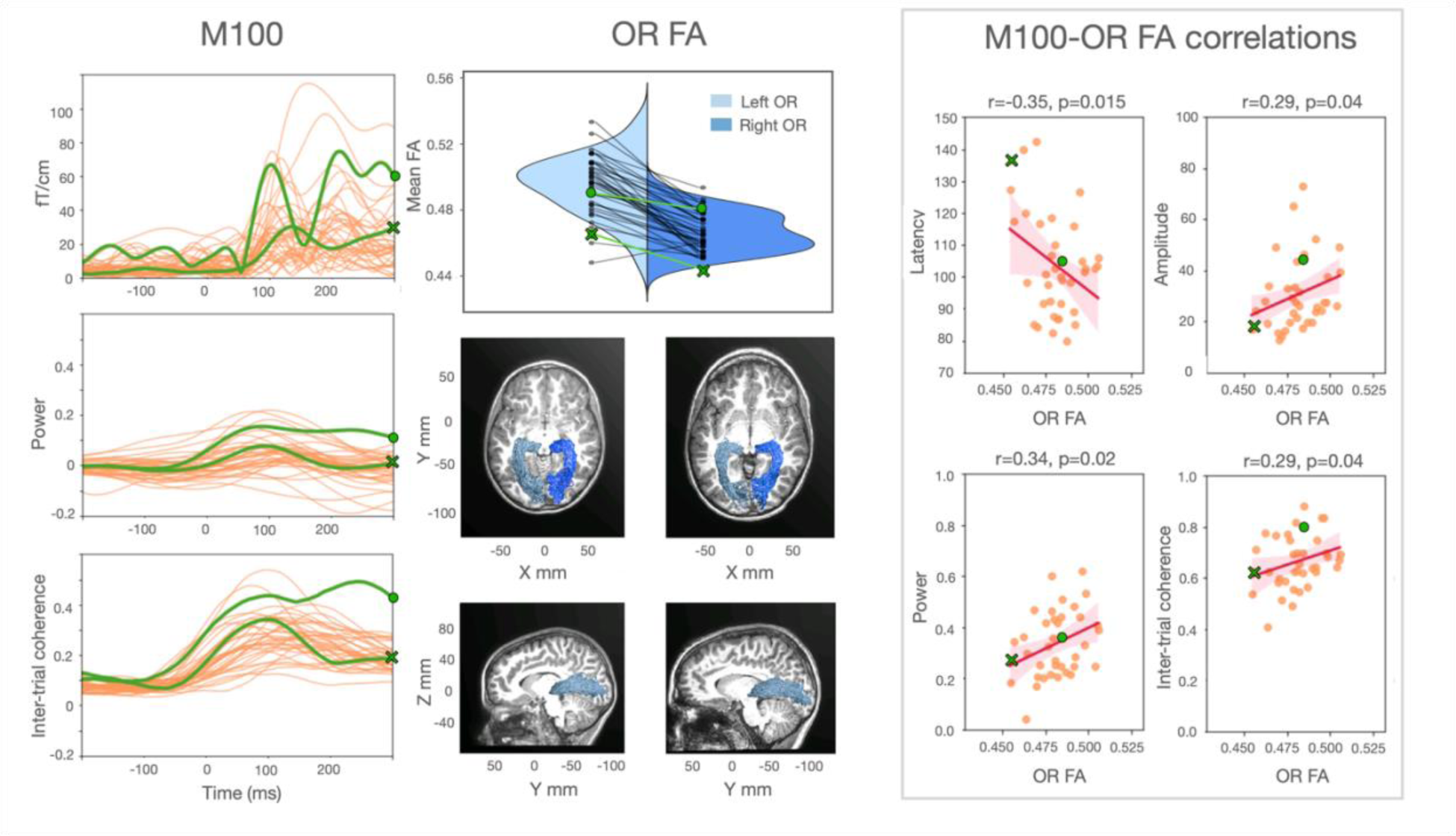
The relation between structural and functional properties of the visual pathways. **M100 panel:** Individual waveforms to high contrast stimuli from a central occipital sensor pair. ERFs, power, and inter-trial coherence average values (between 1 and 8 Hz) are displayed over time. The occipital responses of two representative subjects (S210: female, 7 years old; and S227: male, 11 years old) are marked in green to highlight individual differences. The dot and the cross green markers correspond to the faster and the slower individual, respectively. **OR FA panel:** Violin plot of the individual FA values from the left and right optic radiations. Green markers correspond to the FA values of the same two representative subjects. Sagittal and axial views of the optic radiations for the two representative participants are overlaid on each subject’s structural image. **M100-OR FA correlations panel:** Correlations between MEG measures and optic radiations FA mean values. Shaded areas represent 95% confidence intervals for the regression estimate, which is estimated through bootstrapping. Data points of the two representative participants are marked with a green dot and a green cross. Low contrast stimuli showed similar, although weaker, findings for amplitude, power, and inter-trial coherence (see Supplementary Materials S2).

FA values of left and right optic radiations were averaged together so that they could be correlated with individual M100 responses (which were also averaged across left and right sensors).

M100 latency correlated with optic radiations FA values (*r*=-0.35, *p*=0.02; robust *r*=- 0.32, *p*=0.03); children with higher FA had faster M100 responses than children with low FA (see Figure 3, right panel). This effect remained significant after accounting for age (*β*=-338, *SE*=193, *t(35)*=1.8, *p*=0.04, adjusted R^2^: 0.12; for a summary of all statistical results see Supplementary Materials S3). Optic radiation mean diffusivity measures did not show a clear relationship with electrophysiology after correcting for age (see Supplementary Materials S4). In addition, the relationship between optic radiations FA values and electrophysiology was not observed for white matter tracts that do not carry visual information from retina to early visual cortex (uncinate: *r*=-0.21, *p*=0.10; corticospinal tract: *r*=-0.10, *p*=0.27; Supplementary Materials S5 and S6 for additional control tracts).

M100 responses and FA values for the optic radiations showed age effects between ages 7 and 12. With greater age, M100 responses were faster (*r*=-0.31, *p*=0.03) and FA values were higher (*r*=+0.30, *p*=0.03, see Figure 4). Optic radiations FA values mediated the effect of age on electrophysiological responses (average causal mediation effect: *β*=- 0.07, *CI* [-0.20; -2.0e-3], *p*=0.04; percentage of age effect that is due to the FA mediator: 28%, *p*=0.02). The effect of age on M100 latency was not significant after adding FA as a mediator (average direct effect: *β*=-0.19, *CI* [-0.42; +0.08], *p*=0.14), indicating that maturational variations of optic radiations FA fully mediated age effects on M100 latency to high contrast stimuli.

**Figure 4.**
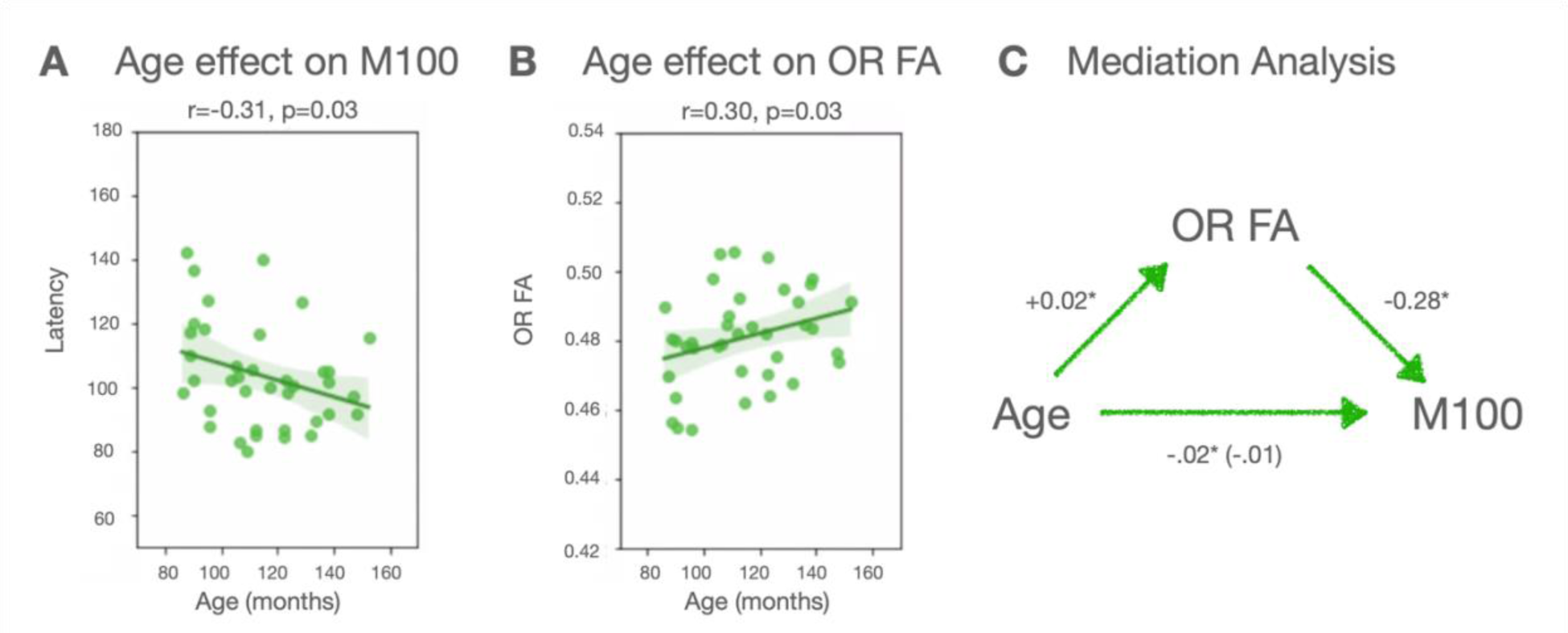
**A and B:** Age effects in M100 latency and optic radiations FA values. **C:** the results of the causal mediation analysis are summarized.

Similar FA correlations were found with the other M100 electrophysiological properties (amplitude: *r*=+0.29, *p*=0.04, robust *r*=+0.35, *p*=0.02; power: *r*=+0.34, *p*=0.02, robust *r*=+0.34, *p*=0.02; inter-trial coherence: *r*=+0.29, *p*=0.04, robust *r*=+0.29, *p*=0.04). Children with high values of optic radiations FA showed higher M100 amplitude, power and inter-trial coherence (see Figure 3 and 4). After correcting for age, the FA effects on M100 amplitude and power remained significant (amplitude: *β*=3.1e-11, *SE*=1.8e-11, *t(35)*=1.8, *p*=0.04, adjusted R^2^: 0.03; power: *β*=3.0, *SE*=1.6, *t(35)*=1.9, *p*=0.03, adjusted R^2^: 0.08; inter-trial coherence: *β*=1.7, *SE*=1.3, *t(35)*=1.3, *p*=0.2, adjusted R^2^: 0.08). Finally, FA mediation effects were confirmed for M100 amplitude and power (amplitude: *β*=6.5e-15, *CI* [+4.2e-17; +1.8e-14], *p*=0.04; power: *β*=6.3e-4, *CI* [+2.4e-5, +1.6e-3], *p*=0.04; inter-trial coherence: *β*=3.6e-4, *CI* [-7.4e-5; +1.1e-3], *p*=0.13).

### Does maturation of the optic tract account for additional variance in electrophysiology?

We next ask whether other stages of the visual pathway additively predict variance in visual responses. The correlation between M100 latency and optic tract FA was not significant (*r*=-0.20, *p*=0.12). Adding the optic tract FA to a regression model including the optic radiations FA did not improve the model fit to the M100 latencies (nested model comparison: *F*(1)=0.13, *p*=.72; adjusted R^2^ of the base model: 0.10; adjusted R^2^ after adding the optic tract: 0.08). Similar results were obtained with the other electrophysiological features (amplitude: *r*=-0.14, *p*=0.20, *F*(1)=3.31, *p*=.08, adjusted R^2^: 0.06, adjusted R^2^ after adding the optic tract: 0.04; power: *r*=+0.14, *p*=0.19, *F*(1)=5e-4, *p*=.98, adjusted R^2^: 0.09, adjusted R^2^ after adding the optic tract: 0.07; inter-trial coherence: *r*=+0.15, *p*=0.19, *F*(1)=0.04, *p*=.85, adjusted R^2^: 0.06, adjusted R^2^ after adding the optic tract: 0.03). Moreover, we did not find a significant correlation between optic tract FA and age (*r*=+.11, *p*=.25). These findings suggest that individual differences in the optic tracts do not account for additional variance in M100 responses.

### Task and stimulus effects

To examine whether the relationship between M100 responses and optic radiations diffusion properties changed as a function of the stimulus or task, we compared different linear mixed effects (LME) models where M100 latency was the dependent variable. We started with a simple model with by-subject random intercepts, including the factors Image Contrast, optic radiations FA, and their interaction. We found a significant main effect of image contrast indicating that low contrast images produce later M100 responses than high contrast images (*β*=5.29, *SE*=0.95, *t(265)*=5.54, p<.001), and a significant FA by contrast interaction indicating that FA only predicted M100 latency for high contrast images (*β*=121.12, *SE*=70.82, *t(264)*=1.71, p=.04; marginal R^2^: 0.10). We progressively added the factors Task, Noise, and their interactions with the other factors. None of these more complex models improved the original model fit (nested model comparison: all χ*^2^*<14, all *ps*>.10; marginal Rs^2^<0.15) and none of the fixed effects reached significance (all *ps*>.06). This was true also for M100 amplitude (all χ^2^<12, all *ps*>.19; marginal Rs^2^<0.23) and power (all χ^2^<12, all *ps*>.17; marginal Rs^2^<0.16). In the models of inter-trial coherence adding the factor Noise and its interactions slightly improved the model fit (all χ^2^(8)=36.6, all *p*<.001; marginal Rs^2^ before and after adding Noise: 0.30 and 0.34), but follow-up analyses just confirmed the main effect of the optic radiations FA (*β*=2.26, *SE*=1.01, *t(36)*=2.25, *p*=.03). Overall, these results do not provide evidence for an effect of stimulus type (e.g., words vs. noise patches) or cognitive task (e.g., color judgment and lexical decision) on the structural-functional link of the visual pathways observed here.

## Discussion

This study describes the link between structural and functional properties of children’s visual pathways and how they change during childhood. We combined dMRI and MEG to measure properties of children’s visual white matter tracts (optic radiations and optic tracts), as well as electrophysiological properties of their visually evoked responses (M100 responses). The data showed that: (1) the structural properties of the optic radiations (indexed by FA values) vary among children and part of this variability is accounted for by age; (2) the electrophysiological properties of the responses in the visual cortex are highly variable in childhood and part of this variability is also explained by age; (3) there is a relationship between the age-related differences observed in visual white matter pathways and those observed in electrophysiology. Specifically, the maturation of the optic radiations during childhood mediates the changes observed in electrophysiological responses of the visual cortex.

These findings complement previous research linking diffusion and electrophysiological properties of the visual pathways in clinical and healthy adult populations (Alshowaeir et al. 2014; Berman et al. 2020; Kolbe et al. 2012; Lobsien et al. 2014; Naismith et al. 2010; M. Y. Takemura et al. 2017; H. Takemura, Yuasa, and Amano 2020). This study not only shows that there is a relationship between structural and functional properties of the visual system, but also that this relationship helps us better understand the development of response properties in the visual cortex during childhood. Between five and twelve years of age the visual system undergoes a large range of transformations, allowing several visual skills to reach their full maturity (e.g., spatial acuity, contrast, and orientation sensitivity; Garey 1984; Siu and Murphy 2018). Part of these developmental differences are reflected by a greater structural coherence and myelination of the visual pathways (with a consequent increase of FA values) and by faster visually evoked responses (Armstrong, Slaven, and Harding 1991; Barnea-Goraly et al. 2005; Dayan et al. 2015; Onofrj et al. 2001). This study revealed that these two types of maturational variations are interrelated and likely, that there is a directional connection that goes from structural to functional developmental changes. Future longitudinal work can establish the temporal sequence of structural and functional changes in the developing brain.

The relationship between electrophysiology and diffusion properties was evident for FA values, and weaker for MD values. This is in line with a trend in the literature that is consistently reporting FA measures as a correlate of electrophysiological responses (Dubois et al. 2008; Gao et al. 2017; Kemmotsu et al. 2012; Lobsien et al. 2014; Price et al. 2017; Shin et al. 2019; Taddei et al. 2012; H. Takemura, Yuasa, and Amano 2020; Westlye et al. 2009; Whitford et al. 2011). Why these effects are more consistent for FA than other diffusion measures such as MD is not clear. FA changes can reflect variations in a myriad of properties including axonal density, size, spatial organization, myelination, as well as changes in glial cells structural properties (De Santis et al. 2014; Jeurissen et al. 2013). Further research on different diffusion-based measures (e.g., axon diameter, quantitative estimate of T1 relaxation) will increase the biological specificity of our brain tissue estimates and improve our understanding of the relationship between white matter microstructure and electrophysiology (Huber et al. 2019; Horowitz et al. 2015).

We could further characterize the link between brain structure and function by examining different visual tracts, stimuli, and tasks. The diffusion properties of the optic radiations and optic tracts were examined and a difference in their developmental trajectories emerged. Age-related differences in FA were evident for the optic radiations, but not for the optic tracts. This suggests that the structural properties of the optic radiations continue maturing during late childhood (Dayan et al. 2015). The optic tracts seemed to be largely stable within this age range. Another possibility is that the optic tract is more affected by partial volume effects (due to its small size). Thus, the signal to noise ratio might be lower for the optic tracts than for the optic radiations making aging effects more evident in the latter ones. As a consequence, developmental differences in electrophysiology were mainly explained by the variation of optic radiations diffusion properties, and no additional variability was explained by the optic tracts. These findings suggest that not all the visual pathways account for visual signal delays over childhood, and that the optic radiation is the best candidate to account for electrophysiological variability in the visual cortex during this developmental window. However, the differences in developmental trajectories between different visual pathways deserves further examination in a larger sample.

By employing different visual stimuli and tasks we examined the extent to which structural-functional connection observed here could be generalized to distinct experimental conditions. Linear mixed effects models showed that children’s M100 modulations depended on the optic radiations properties, and on the level of image contrast. Other experimental factors such as the level of noise of the image and the type of task performed did not have a significant impact on the M100, or on the M100-FA relationship. This suggests that the diffusion properties of the optic radiations predict a wide range of electrophysiological responses, which can be observed with different visual stimuli (readable and unreadable) and tasks (fixation and lexical decision tasks).

Moreover, this structural-functional connection is particularly evident when high contrast stimuli are employed to evoke a highly reliable M100 response.

The present findings also allowed us to expand the functional significance of white matter diffusion properties by relating them to a number of electrophysiological characteristics beyond latency (amplitude, power inter-trial coherence). Past studies have consistently proposed the latency of visually evoked responses as the most likely functional correlate of white matter structure (Adibpour, Dubois, and Dehaene-Lambertz 2018; Alshowaeir et al. 2014; Berman et al. 2020; Dubois et al. 2008; Kolbe et al. 2012; Lobsien et al. 2014; Naismith et al. 2010; Price et al. 2017; M. Y. Takemura et al. 2017; H. Takemura, Yuasa, and Amano 2020). This choice is based on the assumption that the structural integrity of white matter tracts (and therefore, their related diffusion properties) has a great impact on the conduction velocity of neural signals (Alshowaeir et al. 2014; Berman et al. 2020; Horowitz et al. 2015; Kolbe et al. 2012; Lobsien et al. 2014; Naismith et al. 2010; M. Y. Takemura et al. 2017). Evoked response latency is partially related to conduction velocity at the cellular level (Joynt 1989; Siivola 1980; Simons et al. 2007) as it mainly reflects the timing of synchronized post-synaptic activity from cortical pyramidal cells (Bressler and Ding 2006). However, ERFs latency can be related to a range of phenomena at the neural level. Changes in ERFs latency can be due to changes in speed of signal propagation, as well as to an increase of temporal dispersion of the neural signal (within and across trials) resulting in a less synchronized neural activity. Our results based on time frequency measures seem to suggest that white matter structural variations are related to a wide range of electrophysiological changes, which do not only include evoked response latency, but also the degree of coherence of neural signals. The correlation between time frequency and diffusion properties suggests that structural-functional relationships in the visual system depend on the coherence with which white matter fibers can deliver neural signals (hence, the level of synchrony of neural oscillations recorded on the scalp). This might depend on myelination (as mainly suggested from previous clinical works), as well as other structural properties such as the homogeneity of axonal organization, coherence, spatial orientation and density. Additional research on the relation between white matter diffusion properties and time frequency estimates is needed in order to uncover the mechanisms underpinning the functional-structural relationship observed here. Note that time frequency measures present some advantages as compared to ERFs measures (latency, amplitude). First, they usually show a reduced noise when single trials are analyzed within narrow frequency bands (Herrmann et al. 2014). Second, they better represent the degree of synchronization of oscillatory activity from large populations of neurons (even when it is not phase-locked to the stimulus onset, (David, Kilner, and Friston 2006; Herrmann et al. 2014; Z. Zhang 2019). These peculiarities of time frequency measures might facilitate the detection of correlations with diffusion properties and provide new insights on their functional interpretation. For example, future studies with larger samples could examine the coherence between signals among multiple brain regions in relation to the tissue properties of the tracts that carry these signals.

In summary, these findings suggest that the maturation of visual white matter pathways over childhood accounts for variations in electrophysiological responses of the visual cortex. This structural-functional relationships is specific to the optic radiations and can be observed across different tasks, levels of visual noise, and electrophysiological properties of the visual responses. The present findings are an example of how relating white matter properties to functional aspects of the brain can help us reach a more nuanced understanding of the link between development of brain connectivity and changes in electrophysiology.

## Conflict of interest disclosure

The authors declare no competing financial interests.

## Funding statement

This project has received funding from the European Union’s Horizon 2020 research and innovation programme under the Marie Sklodowska-Curie grant agreement No 837228 and Rita Levi Montalcini to SC and was supported by the National Research Foundation of Korea (NRF) grant funded by the Korea government (MSIT) (No. NRF-2019R1C1C1009383) to SJJ. This work was also supported by NSF/BSF BCS #1551330, NICHD R01HD09586101, NICHD R21HD092771 and Jacobs Foundation Research Fellowship to JDY and RF1MH121868-01 to AR and JDY.

## Data availability statement

Data are released through AFQ-browser https://sendycaffarra.github.io/OpticRadiation

## Ethics approval statement

The study was approved by the Institutional Review Board.

## Supplementary Materials S1 Optic radiations segmentation pipeline

**Figure S1.**
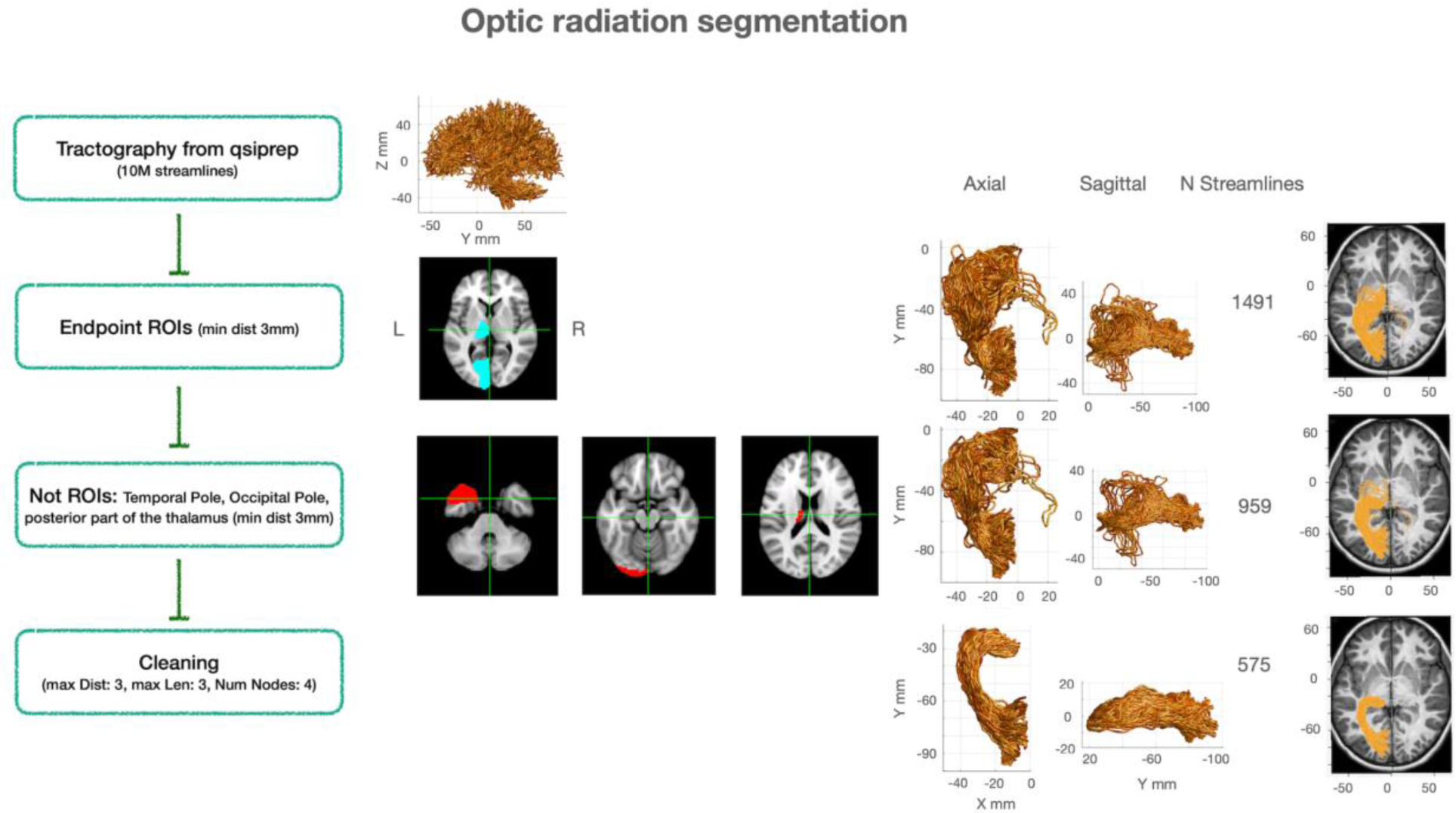
Schematic representation of the optic radiations segmentation. On the left side, each step of the pipeline is described. The middle column shows the tractography reconstructed from qsiprep, and the ROIs used to track the bundle. On the right side, the same optic radiation of an example participant is plotted for each step of analysis.

## Supplementary Materials S2 Low contrast stimuli

**Figure S2.**
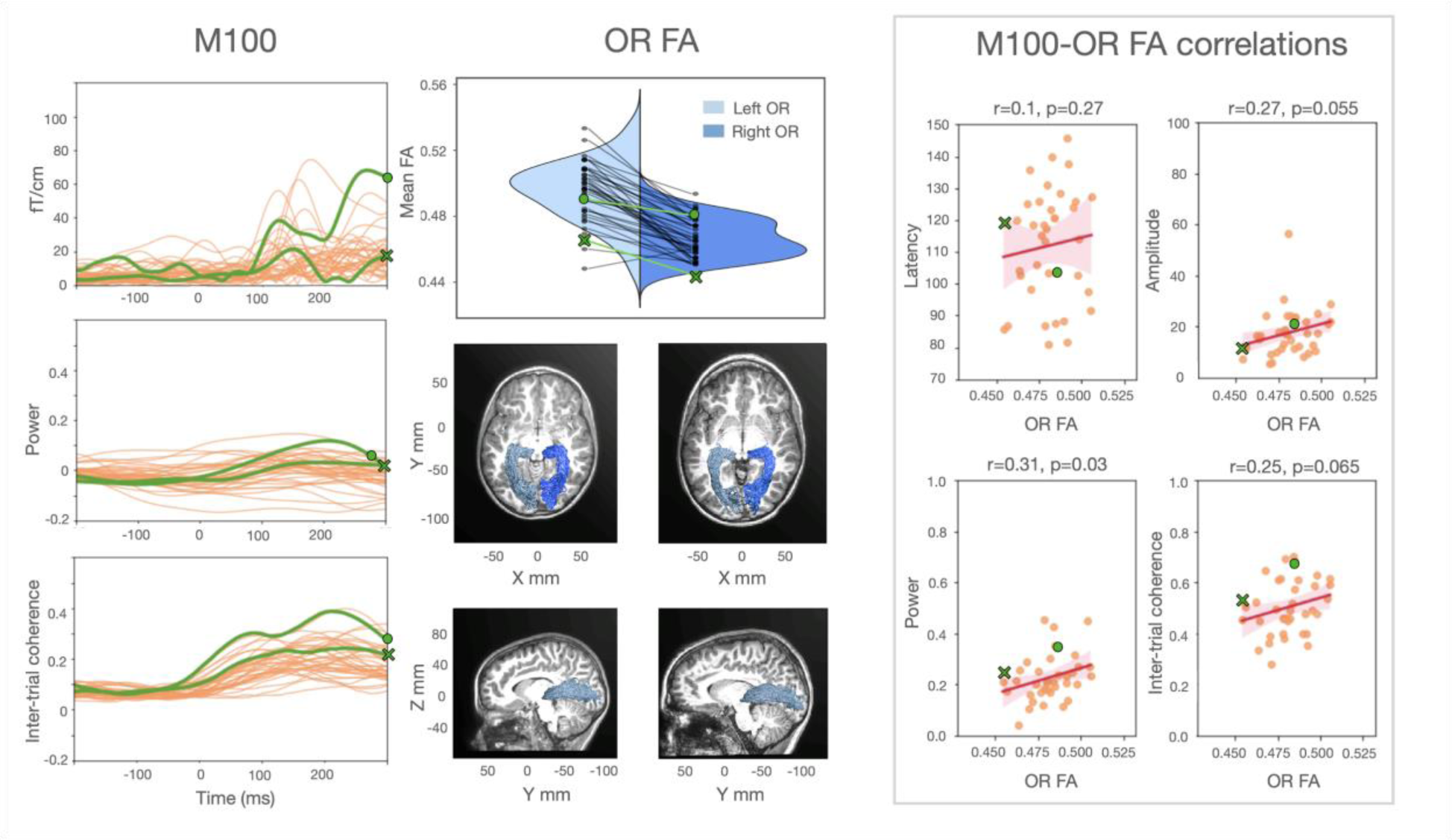
The relation between structural and functional properties of the visual pathways in the low contrast condition. **M100 panel:** Individual ERFs, power and inter-trial coherence average values (between 1 and 8 Hz) are displayed over time. The occipital responses of the same two representative subjects shown in Figure 3 are highlighted (green dot for the faster response and green cross for the slower one). **OR FA panel:** Violin plot of the individual FA values from the left and right optic radiations. Sagittal and axial views of the optic radiations for the two representative participants are overlaid on each subject’s structural image. **M100-OR FA correlations panel:** Correlations between MEG measures and optic radiations FA mean values (robust correlations for latency: *r*=+0.09, *p*=0.30; amplitude: *r*=+0.37, *p*=0.01; power: *r*=+0.31, *p*=0.03; inter-trial coherence: *r*=+0.26, *p*=0.06). Shaded areas represent 95% confidence intervals for the regression estimate, which is estimated through bootstrapping. A full FA mediation effect was visible only for the M100 amplitude (average causal mediation effect: *β*=4.8e-15, *CI* [+1.3e-16; +1.2e-14], *p*=0.04; average direct effect: *β*=-1.0e-14, *CI* [-3.1e-14; +5.7e-15], *p*=0.26)

## Supplementary Materials S3 Summary of statistical results

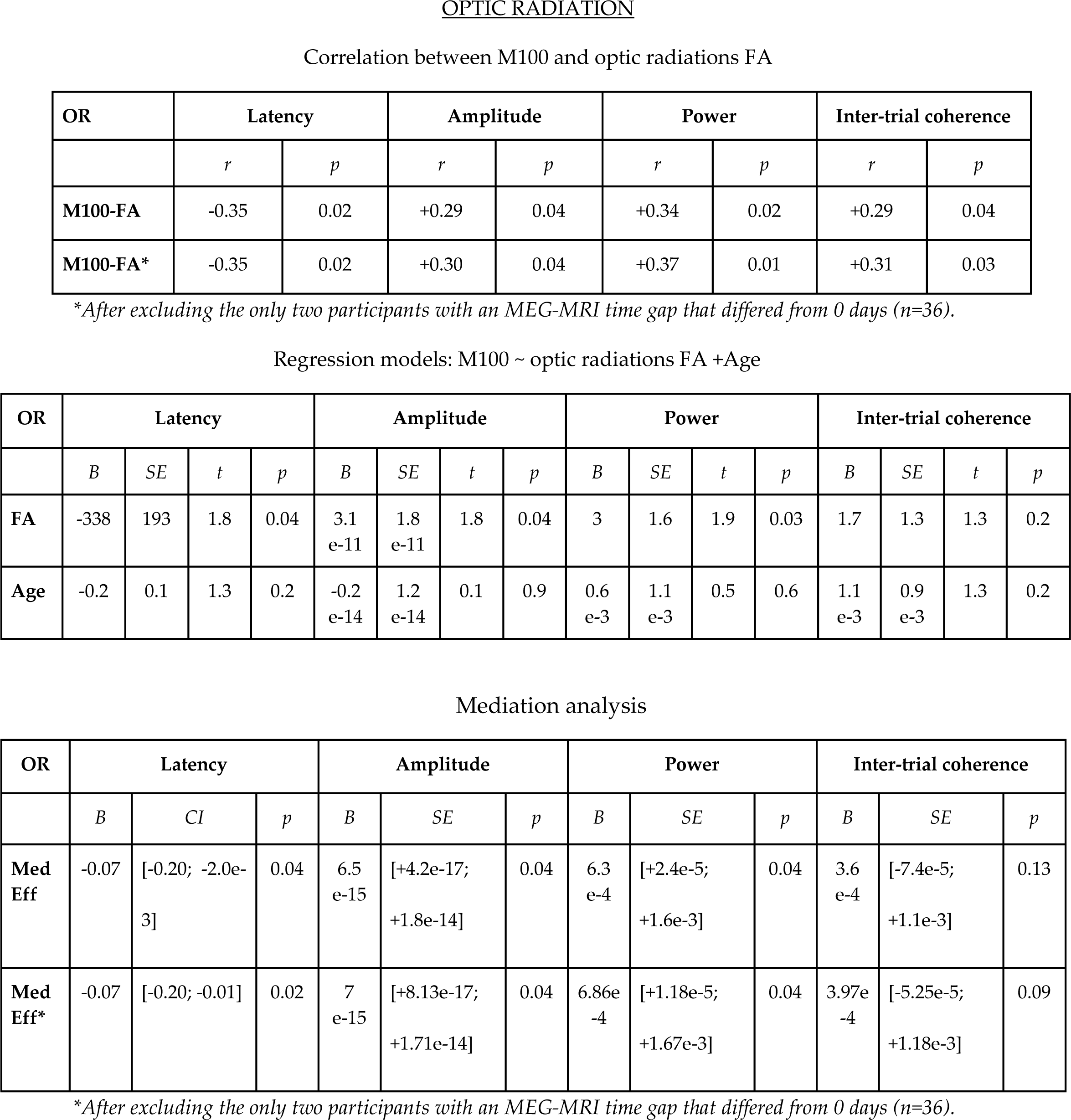
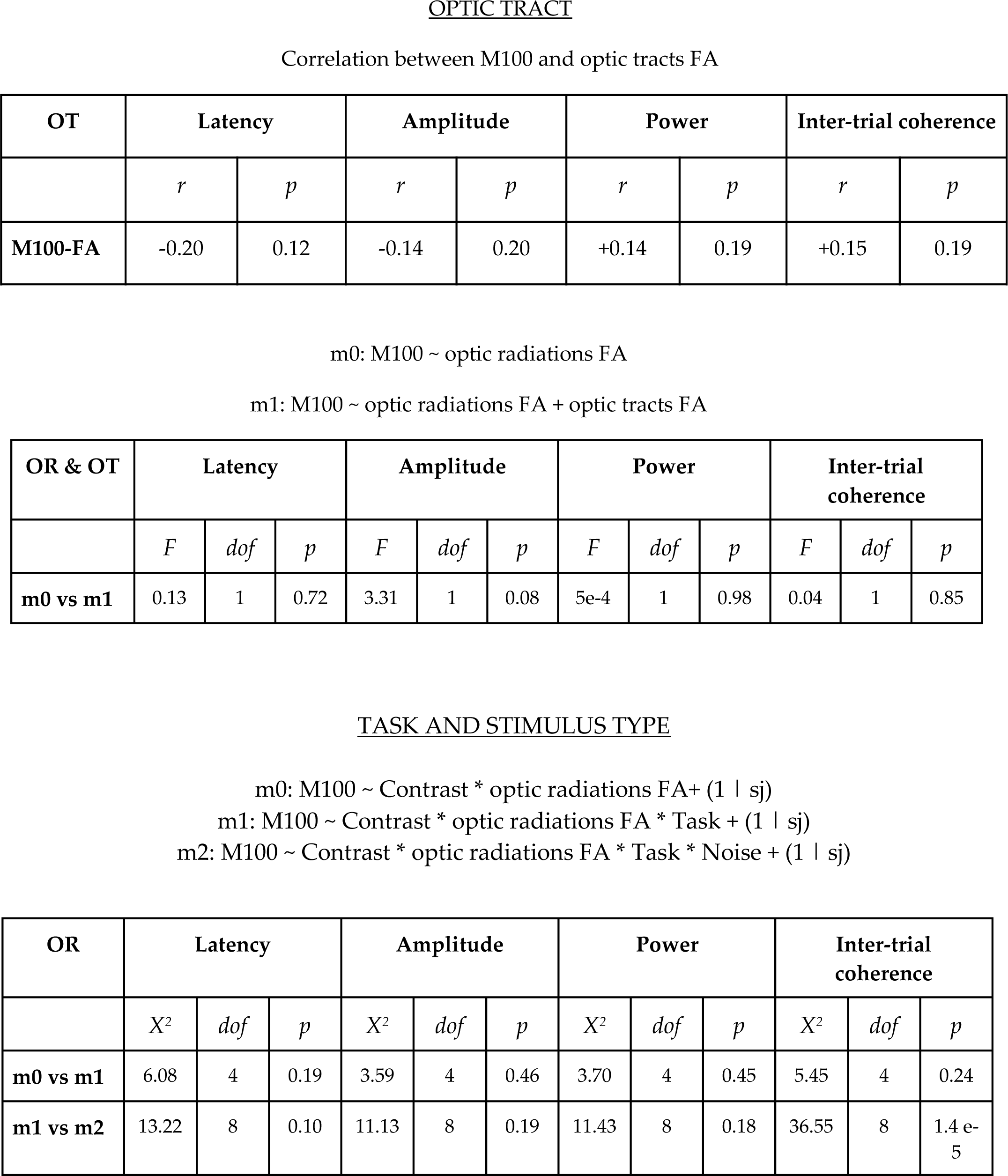

The estimates and the confidence intervals of fixed effects from m2 are plotted here below. The effect of FA on electrophysiological measures does not significantly interact with either Task or Noise.

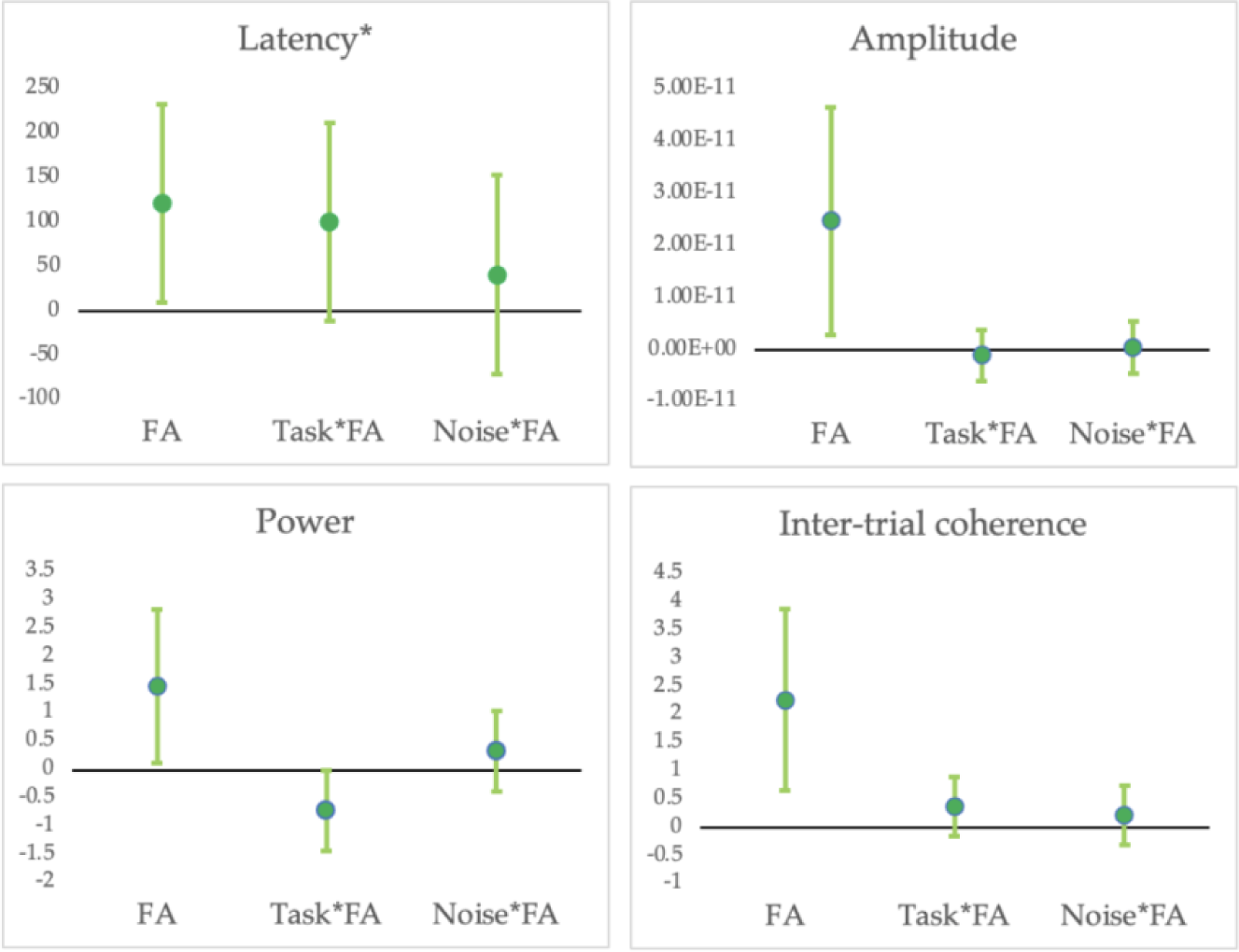

* For Amplitude, Power and Inter-trial coherence the first effect plotted represents the main FA effect (for both high and low contrast stimuli). For latency (*), the interaction between Contrast and FA is plotted because the effect is significant only for high contrast stimuli.

## Supplementary Materials S4 Summary of statistical results for MD

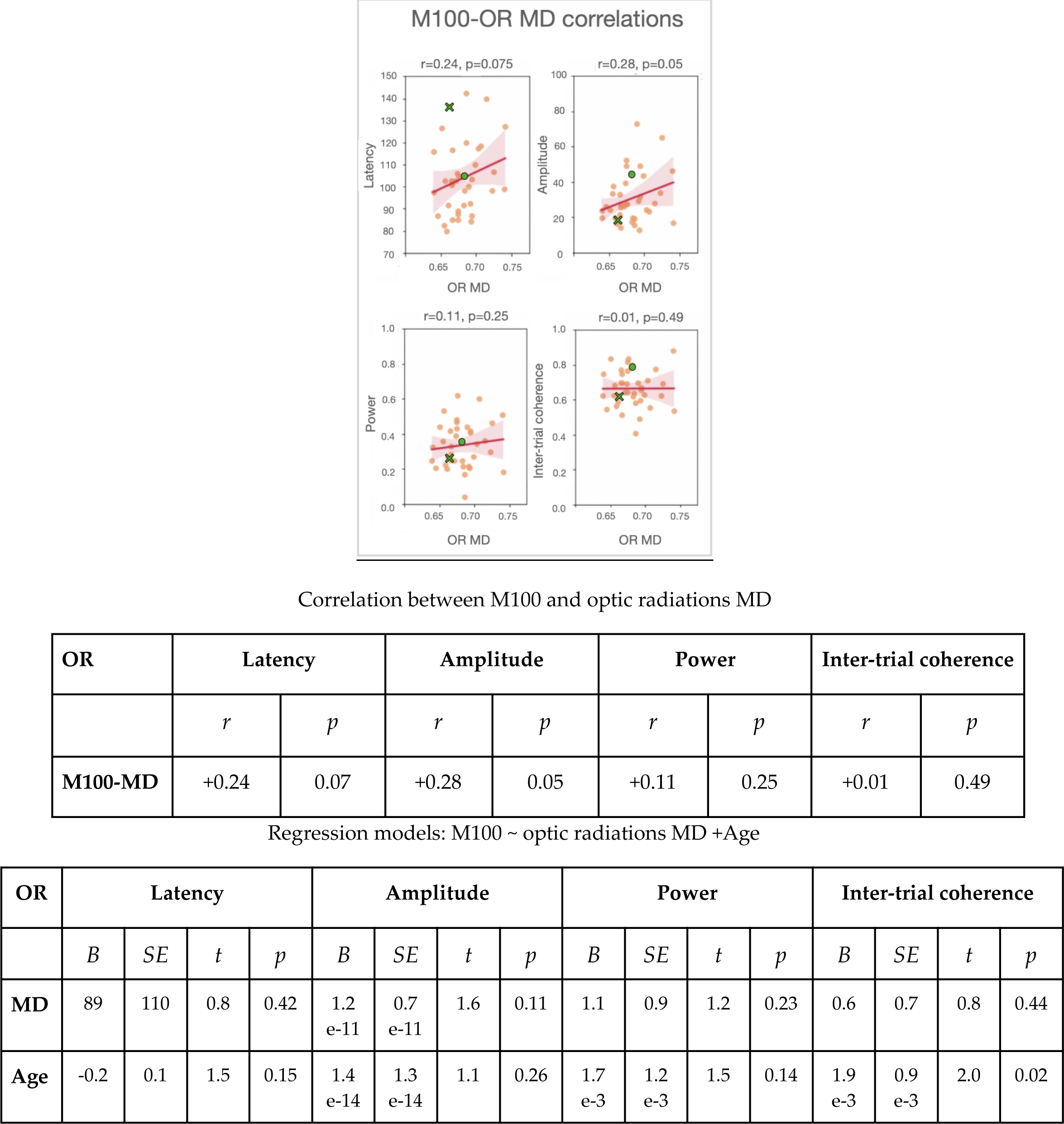

## Supplementary Materials S5 Control tracts

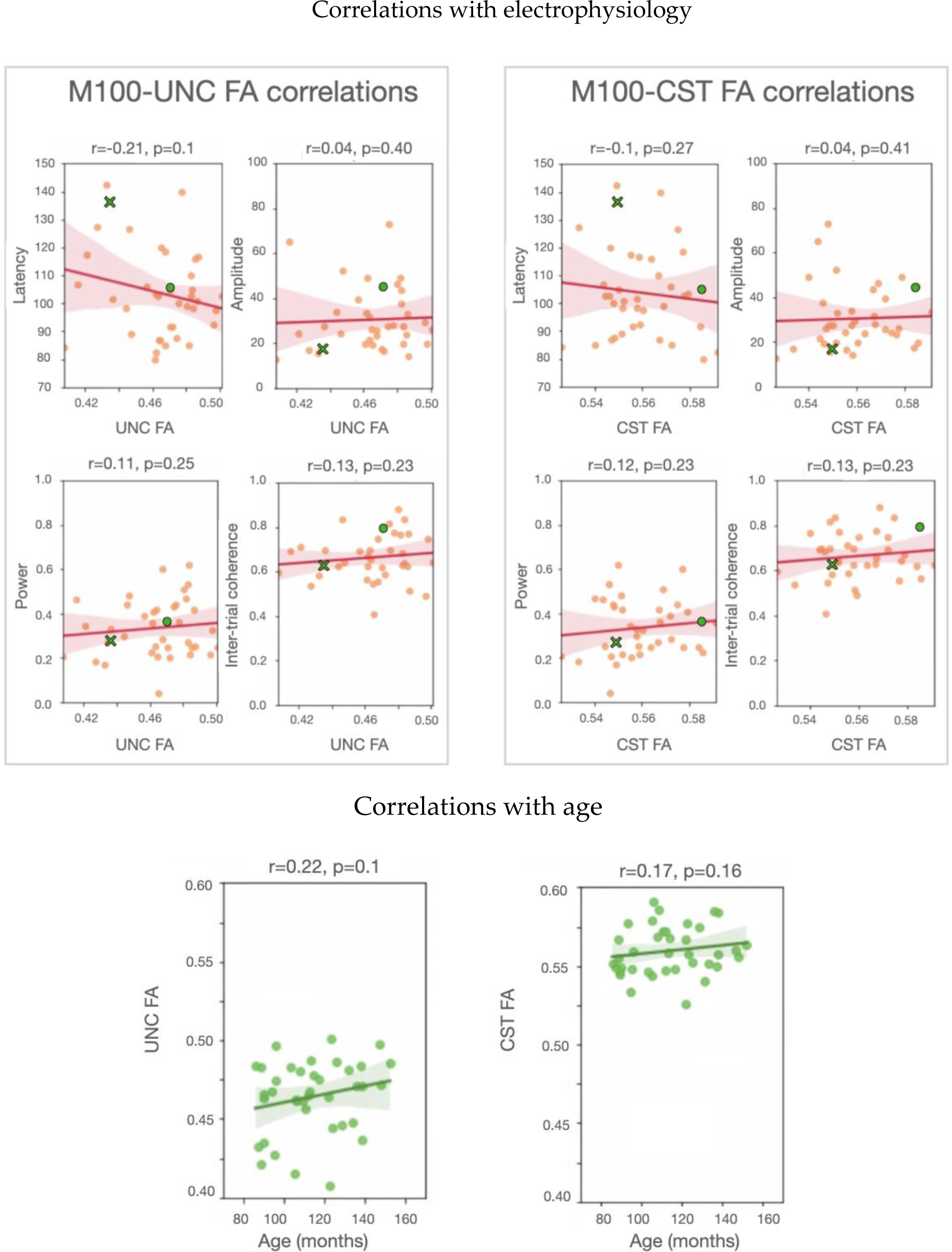

## Supplementary Materials S6 Additional control tracts Correlations between M100 latency and a list of white matter tracts

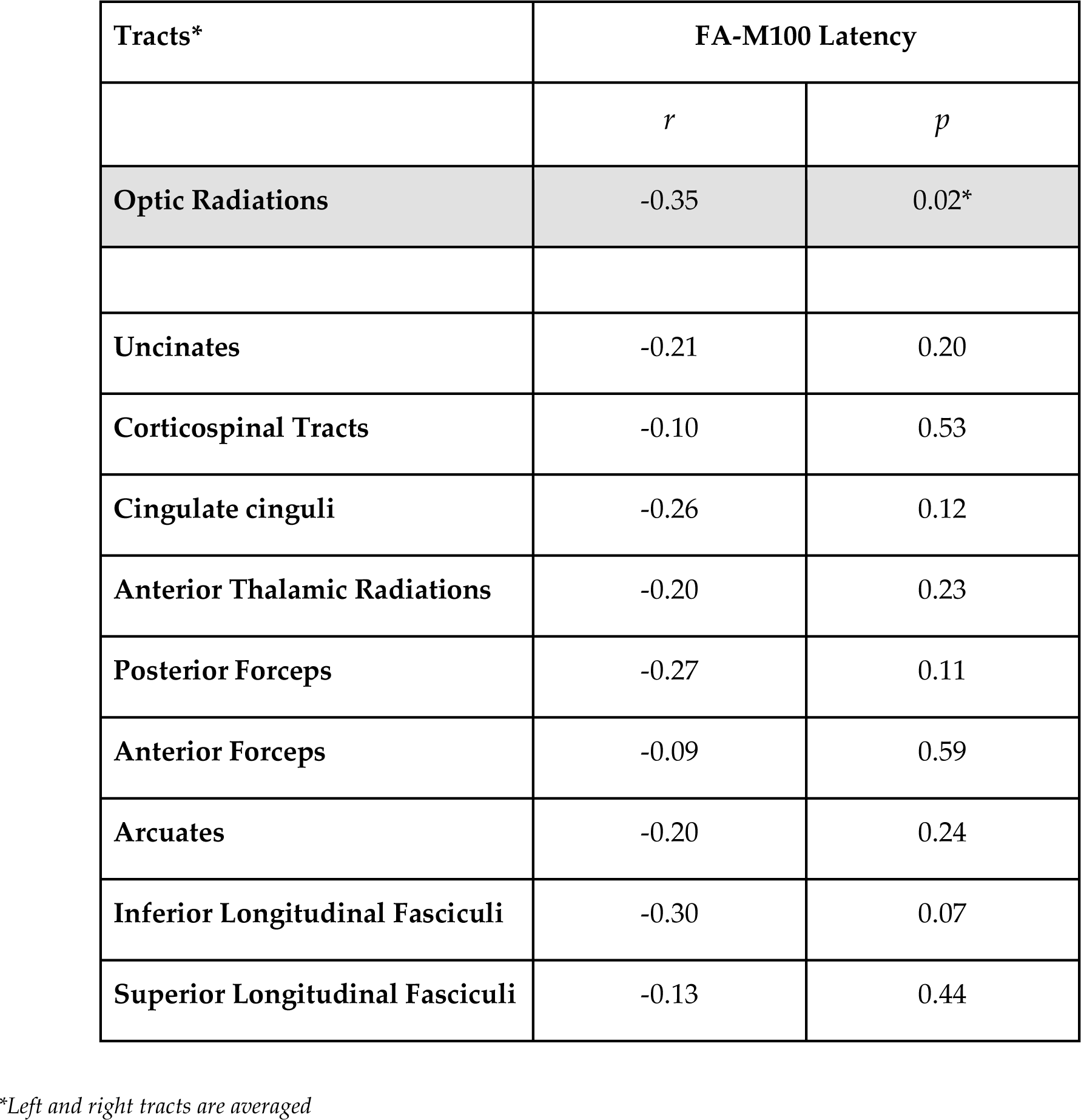

## Notes

### Competing Interest Statement

The authors have declared no competing interest.

